# Evaluation of the efficacy of a full-spectrum medicinal cannabis plant extract with less than 0.3% Δ9-tetrahydrocannabinol in *in vitro* models of inflammation and excitotoxicity

**DOI:** 10.1101/2024.01.10.575133

**Authors:** Emily Ross-Munro, Esra Isikgel, Bobbi Fleiss

**Affiliations:** RMIT University, STEM College, Victoria, Australia; Fenix Innovation Group Pty Ltd, Melbourne, Australia

**Keywords:** Autism, botanical synergy, cytokines, full-spectrum cannabis extract, neurodevelopmental disorders

## Abstract

The rapid development of research on the therapeutic benefits of medicinal cannabis, in parallel with an increased understanding of the endocannabinoid system, has driven research of *Cannabis sativa* constituents for managing neurological conditions. While most studies have focused on the therapeutic potential of the major components of cannabis plant extract isolated or combined, limited research has explored the pharmacological benefits of whole cannabis plant extract. In this study, we investigated the potential anti-inflammatory and neuroprotective effects of NTI-164, a novel full-spectrum cannabis extract with negligible Δ9-tetrahydrocannabinol (THC), compared with cannabidiol (CBD) alone in BV-2 microglial and SHSY-5Y neuronal cells. The inflammation-induced upregulation of microglial inflammatory mediators, being tumour necrosis factor α (TNFα), granulocyte-macrophage colony-stimulating factor (GM-CSF), inducible nitric oxide synthase (iNOS), and Arginase-1 (Arg-1), were significantly attenuated by NTI-164. This immunomodulatory effect was not observed upon treatment with isolated CBD. Compared to CBD alone, NTI-164 prevented elevated mitochondrial activity while normalising cell numbers in immune-activated microglia cells. NTI-164 also promoted the proliferation of undifferentiated neurons and the survival of differentiated neurons under excitotoxic conditions. Overall, our work shows that the anti-inflammatory and neuroprotective effects of NTI-164 as a full-spectrum cannabis extract are enhanced relative to that of CBD alone, highlighting the potential therapeutic efficacy of NTI-164 for the treatment of neuropathologies such as autism spectrum disorder (ASD) and related neuropathologies. This study has further shown that understanding the synergistic effect of phytocannabinoids is integral to realising the therapeutic potential of full-spectrum cannabis extract to inform the design of botanical-derived treatments for managing neurological disorders.

## Introduction

Despite differences in clinical profiles, genetics, and symptoms among neurological diseases, many central nervous system (CNS) disorders share substantial similarities in cerebrovascular impairments and immune dysregulation associated with neuroinflammation as the underlying pathophysiological mechanism of disease [1, 2]. In particular, the role of the neuroimmune axis in the pathogenesis of neurodevelopmental disorders, including autism spectrum disorder (ASD), has been the focus of intense research during the past decade [3–6]. The hallmarks of ASD are social and behavioural deficits, often associated with intellectual disabilities, hyperactivity, anxiety, epilepsy and cognitive impairments [7, 8]. Despite the rising prevalence of this neurodevelopmental disorder [9, 10], significant economic burden [11, 12], and ongoing research efforts, the underlying etiopathogenesis of ASD remains unclear. The aetiology of ASD involves intricate interactions between multiple genetic and environmental risk factors during the critical period of nervous system development, resulting in aberrant neuroplasticity with long-term alterations in brain physiology and behavioural processes [13, 14]. The complex clinical profile of ASD, compounded with significant genetic heterogeneity, has further hindered efforts to dissect underlying disease mechanisms and develop effective pharmacological therapies [15, 16]. Consequently, there are no effective medications to address the core symptoms of ASD, with the current pharmacotherapies targeted at ASD comorbidities frequently associated with adverse side effects and relatively low efficacy [17, 18].

Accumulating evidence highlights the potential role of neuroglial cells and, in particular, disruption of homeostatic microglial function directly contributing to ASD etiopathogenesis [5, 6]. Microglia are myeloid cells of mesodermal origin and the only glial cells to enter the developing brain preceding neurogenesis during the perinatal period [19, 20]. Microglia have a diverse array of physiological functions in adulthood and mediate key developmental processes, including astrocyte formation, early synaptogenesis, synaptic plasticity, synaptic remodelling (pruning) plus the regulation of adult neurogenesis [5, 19, 21, 22]. In response to signs of homeostatic disturbance, microglia shift activity states and transform from homeostatic to an ‘activated state’, such as immune activation, which can result in distinct response phenotypes [23]. Depending on the pathological condition and its impacts on the local microenvironment, microglia contribute various immune-related activities through the differential release of molecules, including pro-inflammatory mediators, anti-inflammatory cytokines, or neurotrophins [5]. The activation states of microglia in response to brain insults have historically been broadly divided into the two reactive phenotypes of an M1 pro-inflammatory response and an M2 neuroprotective response [22]. Although an oversimplification [24], these terms are helpful for basic function descriptions in simplified systems, such as *in vitro* paradigms like those in this study.

Consistent with human and animal studies that have demonstrated glial abnormalities in ASD-related phenotype [3, 13, 25–28], several imaging studies have further reported the presence of markedly immune-activated microglia cells with morphological alterations in multiple brain regions of young ASD patients [29–32]. The role of microglia in neuroinflammation in these individuals is also supported by the observation of differential expression of inflammatory cytokines in different brain regions of children with ASD [33, 34].

Endogenous endocannabinoids, together with their receptors (CB1 and CB2) and associated metabolic enzymes (the endocannabinoid system, ECS), are critical regulators of early neuronal plasticity [35] and behavioural processes [36]. In immune-activated microglia, CB2 receptors modulate synaptic activity and neuroinflammatory responses [37]. A large body of evidence further supports the involvement of ECS in ASD-related neurodevelopmental processes, as aberrant endocannabinoid signalling pathways are linked to ASD pathogenesis [38–40]. Moreover, recent clinical studies in children with ASD have demonstrated altered endocannabinoid-CB2 cellular signalling [41] and circulating endogenous cannabinoids [42, 43], underpinning the importance of the ECS as a potential therapeutic target for ASD and related neuroinflammatory diseases.

The mounting evidence for the involvement of ECS in ASD-related neurodevelopmental processes, in parallel with the well-researched therapeutic potential of medicinal cannabis, has heightened interest in using cannabis-derived compounds to treat neurodevelopmental disorders. For centuries, the therapeutic benefits of *Cannabis sativa* L. extracts have been widely recognised in the treatment of a broad spectrum of nervous system-related conditions ranging from common neurological and hyperexcitability disorders such as epilepsy, affective disorders (e.g. anxiety) to neurodegenerative disorders such as Alzheimer’s disease [44]. The role of the most abundant non-psychoactive phytocannabinoid cannabidiol (CBD) in the modulation of inflammatory and immune responses has been extensively documented [45–47]. Most current research on treating common neurological and psychiatric disorders is focused on the application of CBD alone or combined with the major psychoactive cannabis component Δ9-tetrahydrocannabinol (THC) [48–50]. Several lines of evidence from experimental models suggest that individual phytocannabinoids derived from the *Cannabis sativa* plant modulate microglial functions by blocking excitotoxic-induced activation [51], inhibiting intracellular calcium increase [52] and modulating inflammatory responses [53, 54].

Clinical evidence is emerging that to improve the safety profile and enhance the therapeutic effects of cannabis-based treatments, we should be administering CBD combined with other phytocannabinoids for treating children with neurodevelopmental disorders such as ASD and behavioural-related symptoms [55–58]. Nearly 25 years ago, the term’ entourage effect’ was coined to describe the synergistic contributions of the multitude of endogenous endocannabinoids in cannabis [59]. The benefits of the entourage effect over single isolates have been demonstrated in preclinical models of neurological disorders [60–62] and cancer [63–65]. Increasingly, specific breeding programs are generating cannabis lines with low THC, often known as hemp plants, but with higher levels of other cannabinoids.

Therefore, this study investigated the entourage effect of a full-spectrum medicinal hemp strain cannabis plant extract with only 0.08% THC (NTI-164) on inflammatory and excitotoxic responses in well-established preclinical *in vitro* models.

## Materials and Methods

### Isolation of NTI-164

NTI-164 sativa plant was grown and harvested following the natural process of open pollination. After harvesting, we ground the dried plant using a commercial herb grinder until the particulates were approximately 1mm in size. This was to increase the surface area of the plant for subsequent isolation of plant extract. The ground plant was then mixed at a ratio of 1g to 10 mL with 100% ethanol, and then this mixture was placed on a rocker at 50 rpm for 4 hours. We then aspirated the mixture into a new tube, and the remaining particulates were removed via centrifugation at 300*G* for 10 minutes. We stored the final extract at −20°C until use, and an aliquot was sent for analysis with ultra-high-performance liquid chromatography (U-HPLC) to assess the purity and composition of the final extract.

### Ultra-high-performance liquid chromatography (U-HPLC)

We undertook experiments using an integrated U-HPLC system and a single quadrupole mass spectrometer (MS) detector with an electrospray ionization (ESI) interface. The following ten cannabinoids were combined: CBD, cannabidivarin (CBDV), cannabidiolic acid (CBDA), cannacannabigerol (CBG), tetrahydrocannabivarin (THCV), cannabinol (CBN), Δ9-tetrahydrocannabinol (Δ9-THC), Δ8-tetrahydrocannabinol (Δ8-THC), cannabichromene (CBC). We prepared a mixture to contain 10 parts per million (ppm) of each of the ten cannabinoids in methanol. All solvents were liquid chromatography-MS grade with standards prepared by diluting with 90% mobile phase B and 10% deionized water. Reference standard solutions previously shown to be suitable for generating calibration curves were obtained as pre-dissolved solutions from Novachem, Cerilliant Corporation (TX, USA).

### Microglial BV2 cell culture

The immortalised microglia cell line, BV2 (EP-CL-0493, Elabscience), was cultured in Rosewell Park Memorial Institute (RPMI) 1640 media containing 2mM L-glutamine (Gibco, Cat# 11875) containing 0.1% gentamycin (Thermo, Cat# 15750060) and supplemented with heat-inactivated newborn calf serum (NCS; Gibco, Cat# 26010-074) at a concentration of 10% for expansion and 5% when plated for experiments. All cells were from between passage numbers 5 and 10. Cells were plated at 45,000 cells/mm^2^ and treated 24 hours after plating. Here, cells were treated with 4µL of either phosphate-buffered saline (PBS; Gibco, Cat# 100100) to serve as a control (i.e., unstimulated cells) or interleukin- 1β (IL-1β; 50ng/mL; Miltenyi Biotec, Cat#130-101-682) plus interferon-γ (IFNγ; 20ng/mL; Miltenyi Biotec, Cat# 130-105-785) to induce inflammation (i.e., immune activation). We based the cell culturing and immune activation techniques described here on established protocols from our lab [66–68]. After 1 hour of post-treatment with either PBS or IL-1β+ IFNγ, we applied a 10µL dose of either NTI-164 extract or CBD (1mg/mL), and we undertook the analysis at 24 hours post-treatment. As NTI-164 extract and CBD were dissolved in ethanol for this investigation, all control (PBS-treated) wells also contained a matched ethanol concentration of 0.1%. An independent replicate (n=1) is cells plated and treated from one stock flask per day.

### Neuronal SHSY-5Y cell culture

The immortalised neural precursor cell line, SHSY-5Y (The American Type Culture Collection, Cat# CRL-2266), was used to assess the effect of NTI-164 or CBD treatment on either neuronal differentiation or in response to excitotoxic injury. For expansion, we cultured cells in Dulbecco’s Modified Eagle Medium (DMEM, Gibco, Cat# 11995) that contained 1g/L D-glucose, 584mg/L L-glutamine, 110mg/L sodium pyruvate, and phenol red, further supplemented with 0.1% penicillin-streptomycin (Sigma-Aldrich, Cat# P4458) and 10% NCS. We plated SHSY-5Y cells at 45,000 cells/mm^2^ for experiments.

For work on differentiated cells, we altered the media to contain 1% NCS and daily delivery of 10uM all-trans-retinoic acid (Sigma-Aldrich, Cat#R2625 dissolved in PBS). Specifically, we differentiated the cells for five days with daily half-volume media changes before the excitotoxicity assays. We exposed the SHSY-5Y to 3 mM glutamate (Sigma-Aldrich, Cat# G815) dissolved in PBS, delivered as a 10uL dose. After 1 hour, cells were treated with NTI-164 or CBD as previously described for BV2 experiments, with analysis conducted 24 hours post-treatment. For undifferentiated culture conditions, cells were plated in DMEM as described above. An independent replicate (n=1) is cells plated and treated from one stock flask per day.

### Multiplex Cytokine/Chemokine Assay

In separate sterile 96-well plates, 300uL of 10% bovine serum albumin (BSA) dissolved in PBS (BSA buffer) was added (to prevent any cytokine loss to the plastic), and we incubated plates for 30 minutes at room temperature, then discarded the contents, and air-dried the plates. We then used these protein-blocked plates to collect BV2 culture media at 24 hours following the initiation of the treatment protocol. Media was immediately centrifuged to remove particulate at 300*G* for 10 minutes with supernatant, then collected in an additional BSA-blocked plate and stored at −80°C until use in multiplex experiments. We measured the cytokine and chemokine levels in the media using a Bio-Plex 200 as per the manufacturer’s instructions (Bio-Rad, Cat# M60000007A). Cytokines and chemokines measured included IL-2, IL-10, IL-5, GM-CSF, and TNFα, as per our previous work [69, 70]. We ran all samples in duplicate with data analysed via the Bio-Plex Manager software from BioRad.

### Immunohistochemistry

Plated cells were fixed for 10 minutes with 2% paraformaldehyde (PFA; VWR Chemicals, Cat# 28794.364) in phosphate buffer and washed for 3×5 minutes with PBS. All washes were for 3×5 minutes throughout the protocol. Plates were stored at 4°C in 150uL PBS containing 0.02% sodium azide (Sigma-Aldrich, Cat# S-2002). Immediately before staining, cells were washed and then incubated for 30 minutes with 100uL of BSA buffer containing 0.01% Triton X (Sigma-Aldrich, Cat# X100). Following this, we removed the 75uL of BSA buffer and added 25uL of primary antibody diluted in PBS to obtain the final concentration, as detailed in Table 1. We incubated the plates overnight at 4°C and washed them the following day. Afterwards, 50uL of the corresponding fluorescent secondary antibody (see Table 1) was applied for 1 hour at room temperature and then washed. To visualise cell nuclei, 4’,6-diamiino-2-phenylindole (DAPI; Invitrogen, Cat# D21490) diluted 1:1000 in PBS was applied for 15 minutes at room temperature with plates subsequently washed. For imaging, photomicrographs were taken using the EVOS M5000 (Invitrogen, Cat# AMF5000) in three fields of view per well from duplicate wells and analysed using Fiji 2 to determine the area coverage of each marker. We undertook the staining and analysis per our previous work [71, 72].

**Table 1.**
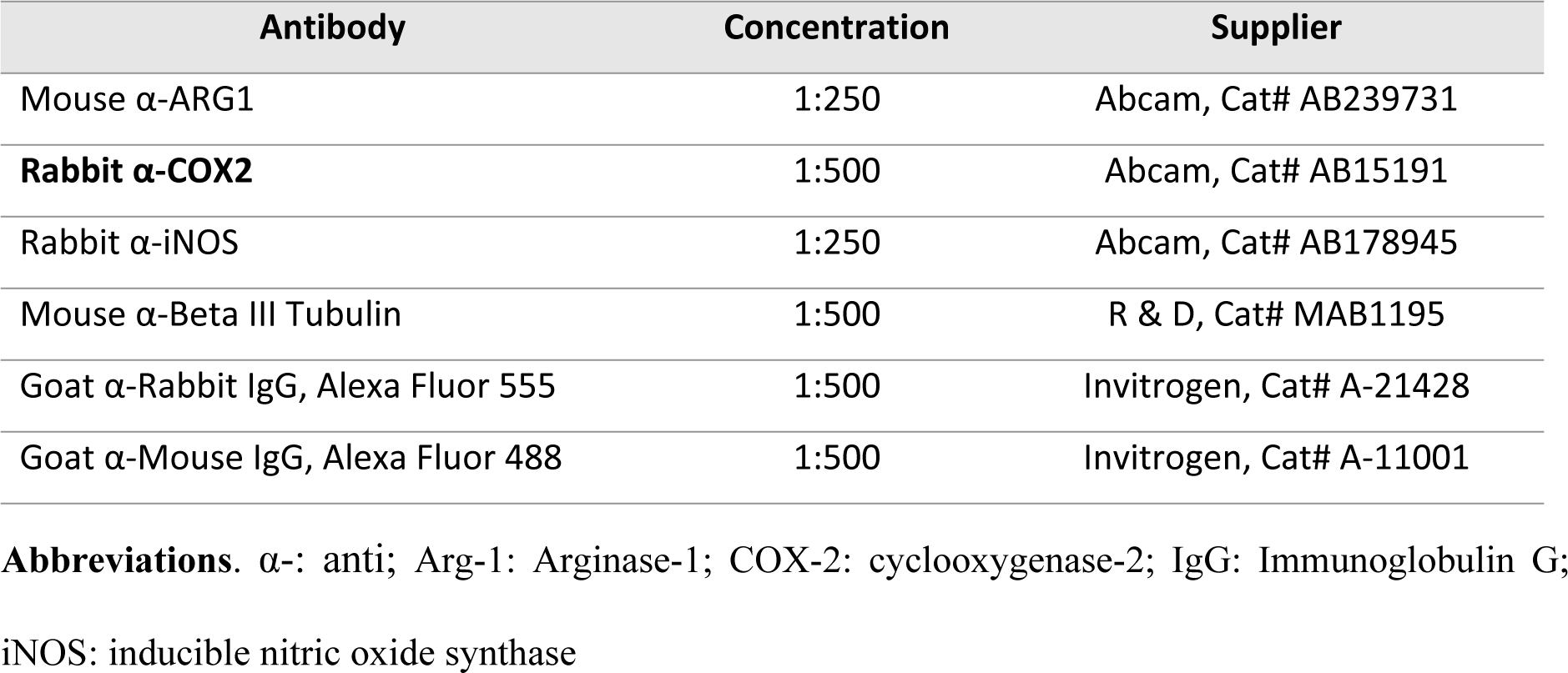
Primary and secondary antibodies.

| Antibody | Concentration | Supplier |
| --- | --- | --- |
| Mouse α-ARG1 | 1:250 | Abcam, Cat# AB239731 |
| **Rabbit α-COX2** | 1:500 | Abcam, Cat# AB15191 |
| Rabbit α-iNOS | 1:250 | Abcam, Cat# AB178945 |
| Mouse α-Beta III Tubulin | 1:500 | R & D, Cat# MAB1195 |
| Goat α-Rabbit IgG, Alexa Fluor 555 | 1:500 | Invitrogen, Cat# A-21428 |
| Goat α-Mouse IgG, Alexa Fluor 488 | 1:500 | Invitrogen, Cat# A-11001 |

**Abbreviations.** $\alpha$ -: anti; Arg-1: Arginase-1; COX-2: cyclooxygenase-2; IgG: Immunoglobulin G; iNOS: inducible nitric oxide synthase

### Cell Viability (Mitochondrial Activity) Assay

We assessed mitochondrial activity using the 3-(4,5-dimethylthiazol-2-yl-)-2,5-diphenyl-2H-tetrazolium bromide (MTT) assay (Sigma-Aldrich, Cat# M6494). As a tetrazolium dye, MTT is bio-reduced by the mitochondria into a formazan product that is insoluble in the tissue culture medium. Upon being dissolved, we quantified the formazan content via a colourimetric analysis. In brief, we added MTT to the wells at a final concentration of 250µg/mL. After 30 minutes, we carefully removed and discarded the media, then dissolved the formazan within the wells in dimethylsulfoxide (DMSO; Sigma-Aldrich, Cat# D2650) and absorbance measured at 490nm using a spectrophotometer (Glomax Multi+; Promega, UK).

### Statistics

We averaged the data for replicates within experiments, and then data from at least three independent experiments was analysed using GraphPad Prism software (GraphPad Software, Inc.). We have outlined the replicate numbers for each experiment and the specifics of the statistics for each analysis in the figure legends. We expressed all data as mean ± standard error of the mean (SEM) with significant differences between groups set at a p-value of less than 0.05 (P<0.05)

## Results

### Chemical characterisation of NTI-164 constituents by U-HPLC

The U-HPLC analysis revealed that NTI-164 was rich in the acidic precursor of CBD, CBDA as the main constituent of the extract (up to 60% wt/wt). The extraction procedure also generated a solution of up to 14% wt/wt CBD, 0.44% wt/wt CBG, 0.06% CBDV and 0.08% wt/wt THC content. The HPLC chromatogram was recorded at 229 nm, and we have shown an example in Figure 1.

**Figure 1.**
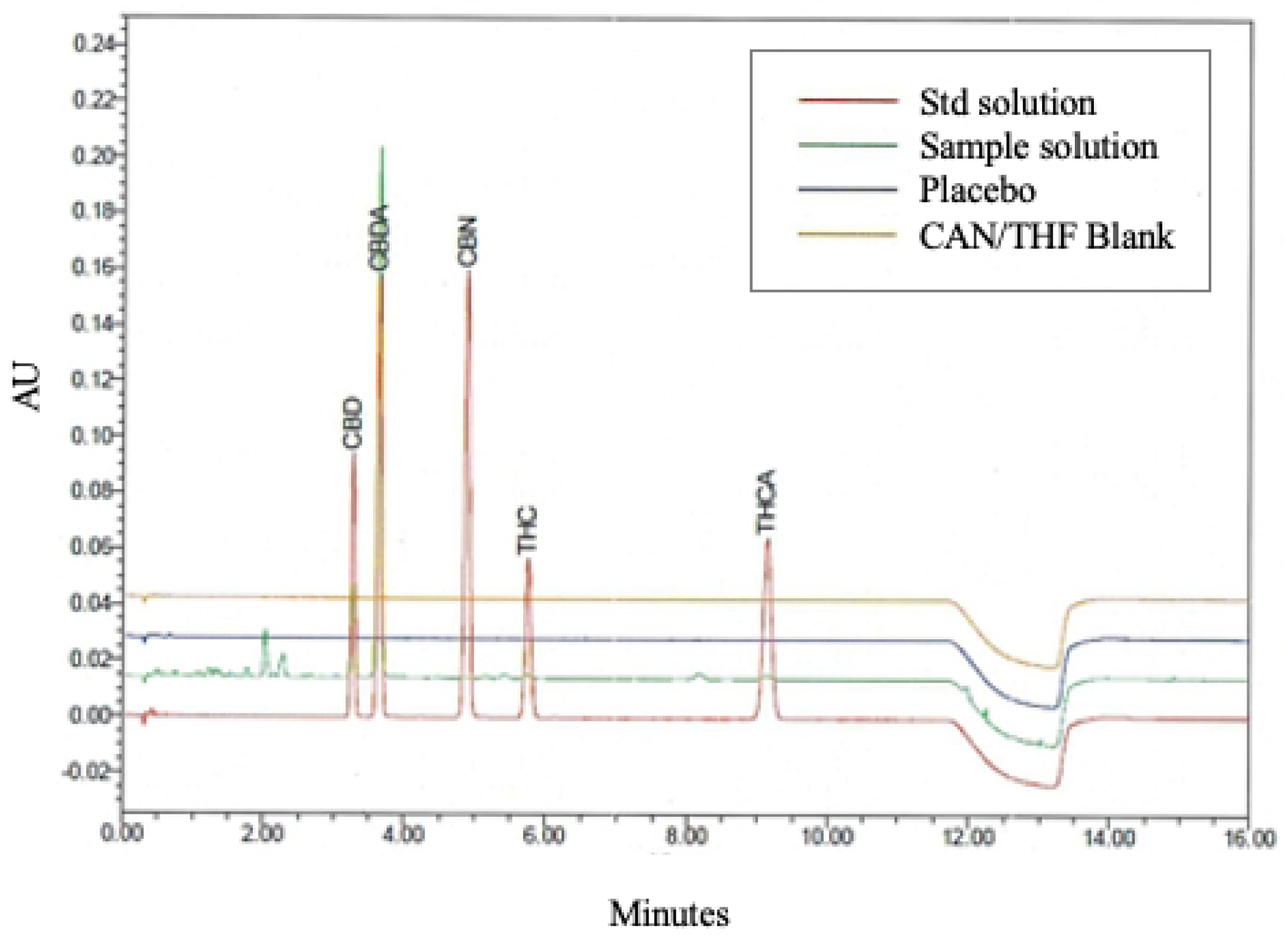
A representative HPLC chromatogram of the NTI-164 extract was recorded at 229 nm, according to the method used to analyse cannabinoids. Representing the main component, the CBD acidic precursor, CBDA eluted at 3.41 minutes, CBD eluted at 3.08 minutes, CBN at 4.57 minutes, THC at 5.35 minutes and THCA at 8.45 minutes. Abbreviations. CBD: cannabidiol; CBDA: Cannabidiolic acid; CBN: cannabinol, THCA: tetrahydrocannabinolic acid; THC:Δ9-tetrahydrocannabinol

### Effect of NTI-164 on energy production and number of BV-2 microglia

In unstimulated BV2 microglial cells (treated with PBS + EtOH excipient), NTI-164 or CBD treatment had no significant effect on mitochondrial activity as assessed via the MTT assay (Figure 2A). In line with prior findings, we observed that mitochondrial activity was significantly increased in immune-activated cells stimulated with IL-1B+IFN-y (+EtOH excipient). While NTI-164 prevented this increase, CBD was not (Figure 2B). In line with this, exposure to NTI-164 or CBD in unstimulated cells did not significantly alter overall cell number as assessed via area-coverage analysis of DAPI staining (Figure 2C). Interestingly, CBD significantly increased the area coverage of DAPI staining in immune-activated cells. We did not observe any impacts of NTI-164 treatment on DAPI staining in the immune-activated cells (Figure 2D).

**Figure 2.**
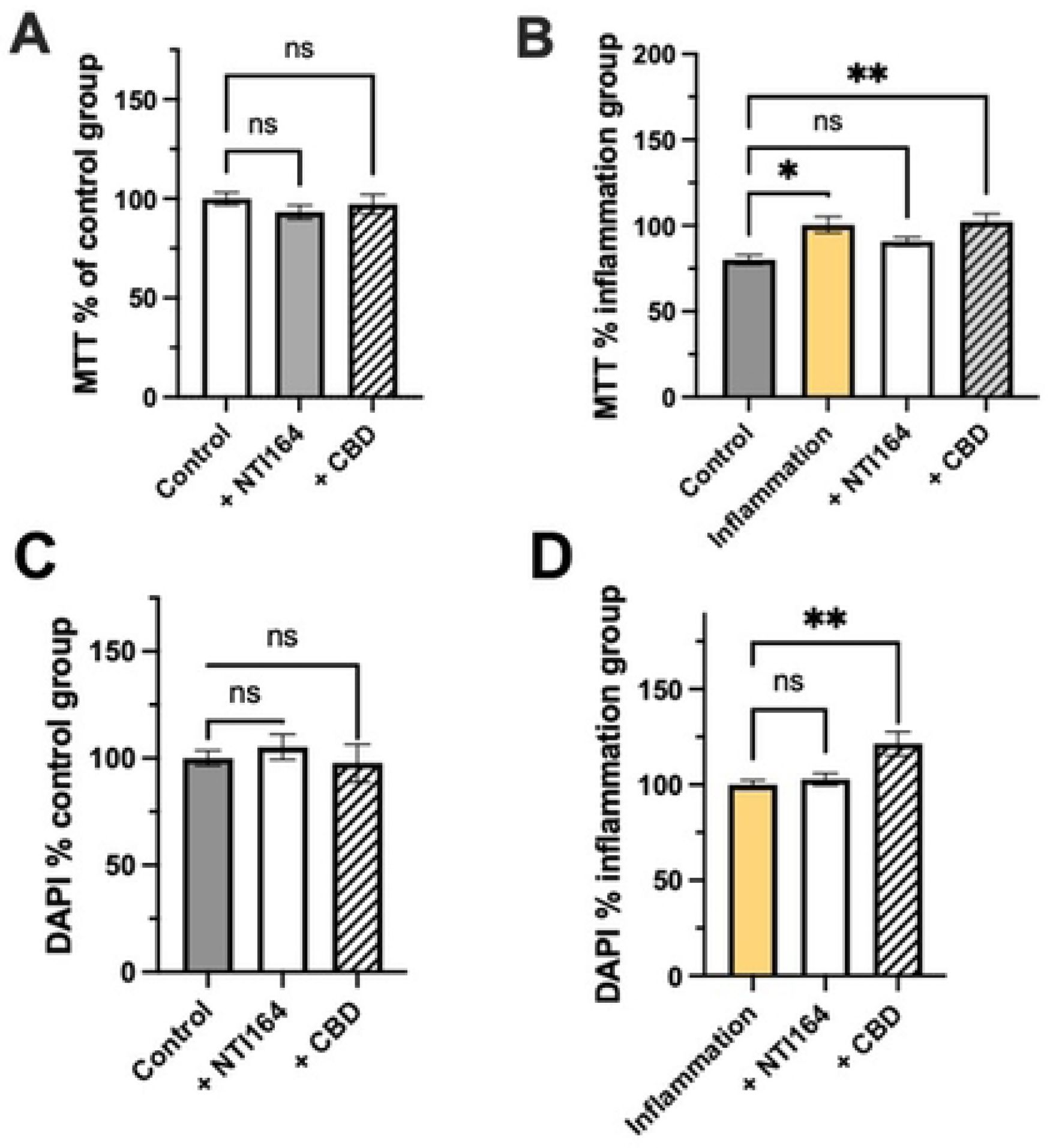
Effect of NTI-164 and CBD on MTT outputs in BV-2 microglia in **A)** unstimulated condition, **B)** immune-activated condition and overall DAPI-positive cell number in **C)** unstimulated condition, **D)** immune-activated condition. Data expressed as Mean ± SEM, n=6-10. Significance set at *, p<0.05; **, p<0.01; ***, p<0.001.

### Effect of NTI-164 on the expression of inflammatory markers in BV-2 microglial

In immune-activated microglia, NTI-164 treatment resulted in a significant decrease in the expression of inducible nitric oxide synthase and arginase-1, which was not observed in CBD-treated cells (Figure 3A, B). Furthermore, NTI-164 and CBD were both able to significantly reduce the expression of COX-2 from the immune-activated microglia (Figure 3C).

**Figure 3.**
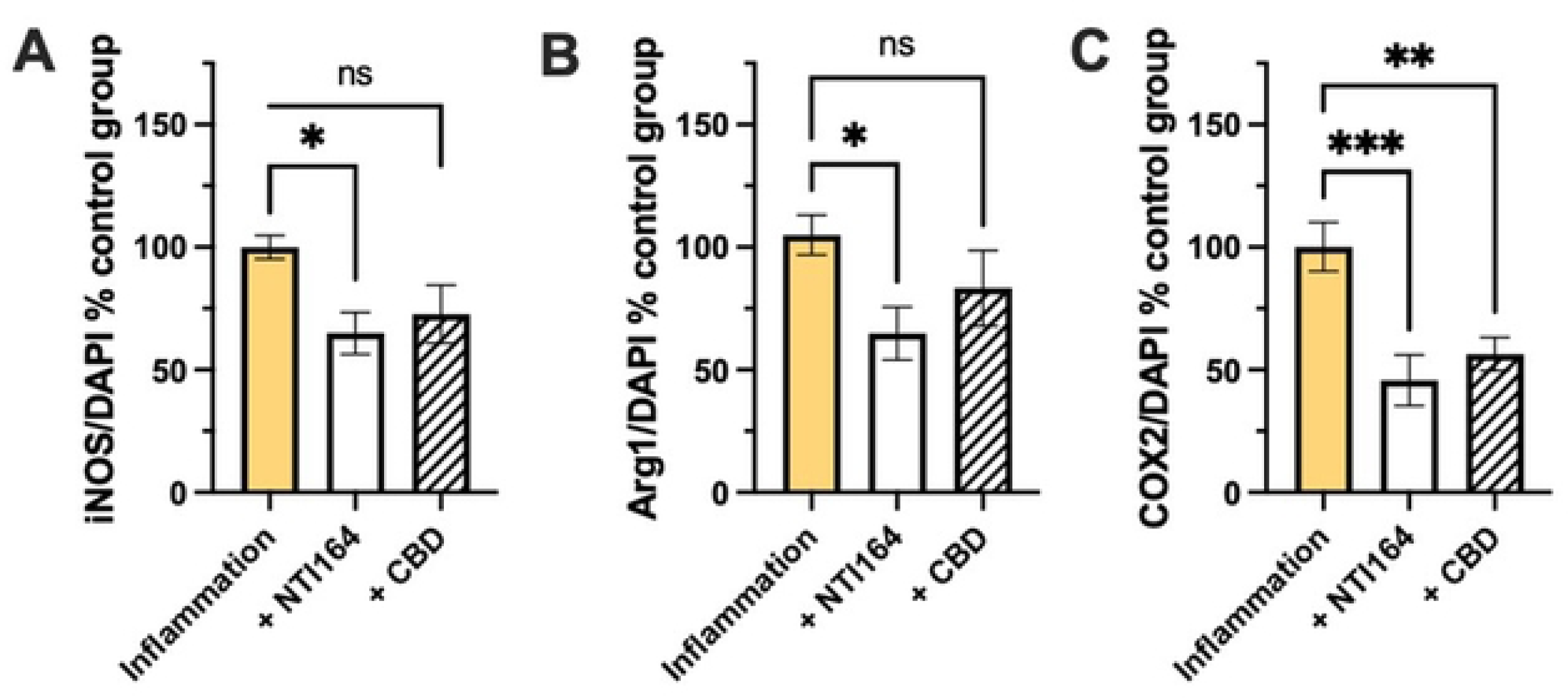
Effect of NTI-164 and CBD on the expression of inflammatory markers in BV-2 microglia cells. **A)** inducible-nitric oxide synthase (iNOS), **B)** arginase-1 (Arg1), and **C)** cyclo-oxygenase-2 (COX2) protein expression normalised to DAPI-positive cell number in the control (PBS only) group. Data expressed as Mean ± SEM, n=6-8. Significance set at *, p<0.05; **, p<0.01; ***, p<0.001.

### Effects of NTI-164 on microglia cytokine production

In immune-activated BV2 cells, treatment with NTI-164 or CBD led to no significant change in the levels of the anti-inflammatory cytokines IL-4 and IL-10 (Figure 4A, B). However, treatment with NTI-164 significantly reduced the expression of the growth and differentiation-inducing chemokine GM-CSF, but this finding was not observed in CBD-treated cells (Figure 4C). NTI-164 and CBD significantly decreased the production of the pro-inflammatory IL-2 (Figure 4D), with no significant alteration to levels of inflammatory cytokine IL-5. Finally, NTI-164 but not CBD was able to reduce levels of TNFα significantly (Figure 4F).

**Figure 4.**
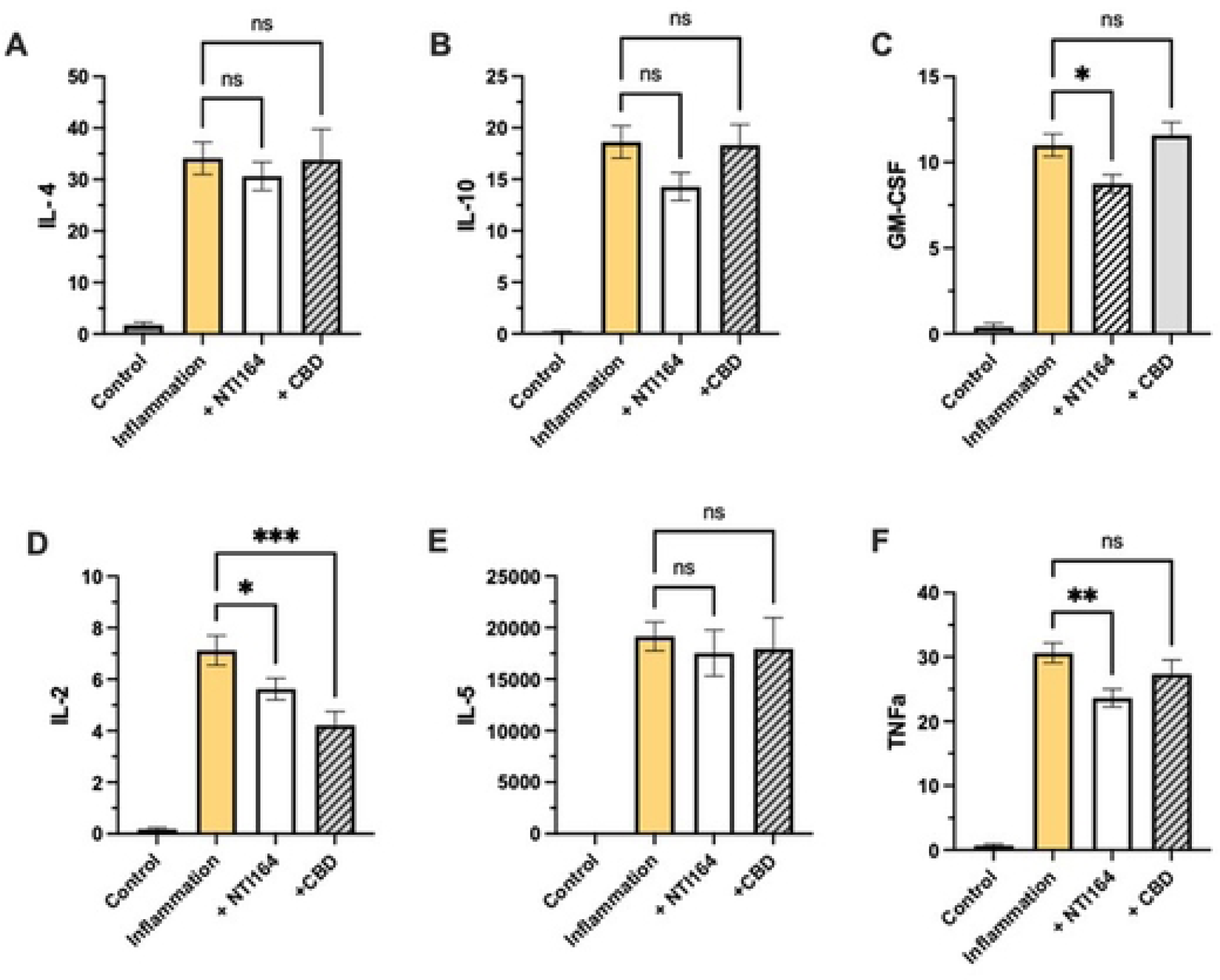
Impact of NTI-164 and CBD on inflammation-induced cytokine release from BV2 microglia. In **A)** interleukin-4 (IL-4), **B)** interleukin-10 (IL-10), **C)** Granulocyte-macrophage colony-stimulating factor (GM-CSF), **D)** interleukin-2 (IL-2), **E)** interleukin-5 (IL-5) and **F)** tumour necrosis factor-alpha (TNF-α) analysed by multiplex immunoassay and expressed as fluorescent units. Mean ± SEM, n=6-7. Significance set at *, p<0.05; **, p<0.01; ***, p<0.001.

### Effect of NTI-164 and CBD on neuronal responses

In undifferentiated neurons, NTI-164 treatment significantly increased the area coverage of DAPI staining, while CBD treatment resulted in no detectable change to DAPI staining (Figure 5A). However, neither treatment induced spontaneous differentiation as assessed via expression of Beta III Tubulin (Figure 5B). In differentiated neurons, neither NTI-164 nor CBD altered the number of cells or the amount of de-differentiation (Figure 5 C, D). In a paradigm of excitotoxicity, NTI-164 significantly increased the mitochondrial activity (survival) of neurons, while CBD could not rescue the mitochondrial activity (Figure 5E).

**Figure 5.**
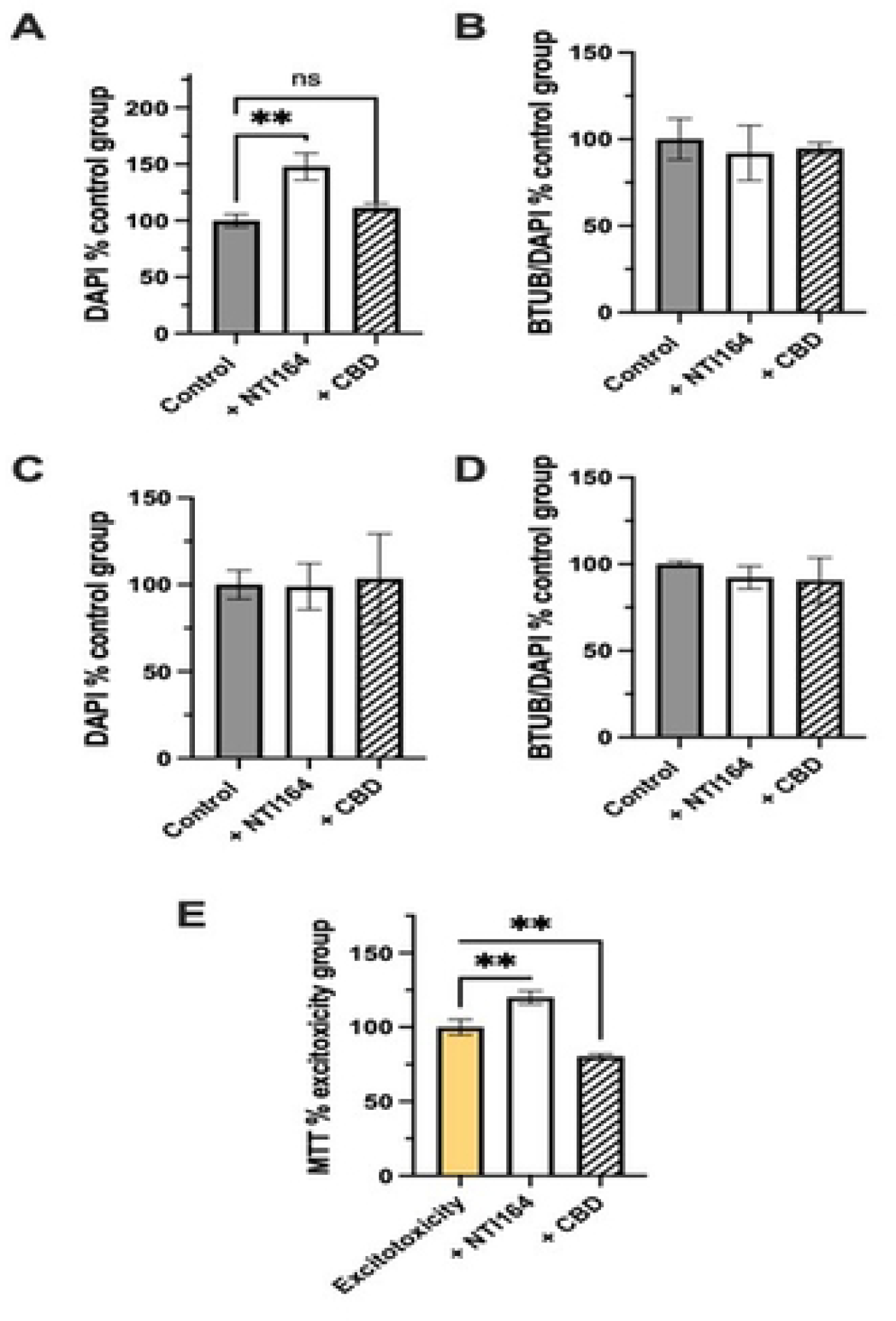
Impacts of NTI-164 and CBD on SHSY-5Y neurons. In **A)** impacts on numbers of DAPI-positive cells in undifferentiated cultures, and in **B)** impacts on the expression of beta-tubulin (BTUB) as a proposition of DAPI-positive cells as an indicator of spontaneous differentiation in undifferentiated cells. In **C)** impacts on numbers of DAPI-positive cells in differentiated cultures, and in **D)** impacts on BTUB expression in differentiated cells, and **E)** impacts on the MTT outputs (mitochondrial activity assay) in differentiated cells in the glutamate exposure excitotoxicity assay. Mean ± SEM, n=6-10. Significance set at *, p<0.05; **, p<0.01; ***, p<0.001.

## Discussion

The current study sought to determine the impact of a full-spectrum medicinal cannabis plant extract (NTI-164) using *in vitro* models of neuroinflammation and neuroinjury. In our study, NTI-164 normalised inflammation-induced changes in immune-activated microglia cells by preventing increased mitochondrial activity and promoting the survival of differentiated neurons under excitotoxic conditions. As a common feature of neurodegenerative and neurodevelopmental disorders, neuroinflammation typically involves activation of resident immune cells of CNS (glia cells) and chronic production of cytokines together with reactive oxygen intermediates resulting in immune cell infiltration, excitotoxicity with subsequent damage cells and to the blood-brain-barrier integrity [73]. In particular, consistent evidence supports the contribution of microglia-mediated neuroinflammatory response in the pathogenesis of ASD [74]. In children with ASD, an abnormal cytokine profile has been reported [75], with elevated levels of pro-inflammatory cytokines predominantly in children with more aberrant behaviours [76]. In parallel, the ECS is involved in modulating microglial phenotype and mediating inflammatory signalling pathways in neuropathology [77]. Compared to neurotypical children, an abnormal profile of ECS has been reported in the peripheral blood of children with ASD [41–43]

Phytocannabinoids or exogenous cannabinoids are abundant in the *Cannabis sativa* plant and have been found to influence microglia regulation of inflammatory response through various biological mechanisms [51, 52]. The therapeutic benefits of combined phytocannabinoids have been consistently demonstrated in several experimental models [52, 78–81] and clinical research studies [82], in particular for the treatment of ASD-related symptoms [56, 57, 83–85]. In an amyotrophic lateral sclerosis animal model, daily treatment of mice with a phytocannabinoid-enriched botanical extract (Sativex) for 20 weeks resulted in a significant decrease in the progression of neurological impairment [81]. In another multiple sclerosis model, treatment with Sativex for ten days reduced microglial immune-related activity and expression of pro-inflammatory cytokines [80]. In a more recent experimental study, THC and CBD combination treatment (but not either compound alone) in a murine multiple sclerosis model led to reduced levels of pro-inflammatory cytokines such as IL-6 and TNF-ɑ while increasing production of anti-inflammatory cytokines [62]. The authors proposed the underlying anti-inflammatory and neuroprotective mechanism of the combined phytocannabinoids to involve cell cycle arrest, apoptosis and induction of anti-inflammatory cytokines in immune-activated brain cells through regulation of miRNA-mediated signalling pathways [62].

In this study, the full-spectrum cannabis extract NTI-164 effectively attenuated elevated mitochondria activity and normalising cell numbers in immune-activated microglia cells and CBD alone did not. We also found that NTI-164 was able to prevent the increase of inflammatory mediators TNFα, GM-CSF, iNOS and Arg-1 and significantly reduced their levels under inflammation-induced conditions. Microglial activation is an important neuroglial response to injury or any pathological event, as microglia are the chief cells engaged in immune surveillance [86]. Animal models and human studies consistently demonstrate an abnormal microglial function associated with ASD-related symptoms. In an inflammatory animal model, microglia activation resulted in ASD-like neurobehavioral deficit [87], which has been linked to impaired ECS signalling, including elevated levels of endocannabinoids and its metabolic enzymes in the amygdala region of the brain [88]. In children with ASD, studies have shown activation of microglia and altered inflammatory gene expressions in multiple brain regions, providing strong evidence supporting that neuroinflammation is involved in ASD pathogenesis [33, 34, 89]. Evidence from a recent systematic review which revealed morphological alterations of activated microglia in various brain regions of individuals with ASD provides additional support for microglia-mediated neuroinflammation in the development of ASD [74]. Disruption in microglia function mediated by neuroactive substances can lead to marked neuronal impairment, contributing to neuronal dysfunction in neurodevelopmental disorders such as ASD [90]. Cannabis extracts may provide potential therapeutic benefits by targeting microglia-induced neuroinflammation in neuropathology.

Both NTI-164 and CBD significantly decreased IL-2 and COX-2 levels in immune-activated cells. Similarly, previous studies have shown that whole cannabis extract [91] and the acidic cannabidiol CBDA [92] exhibit anti-inflammatory effects by inhibiting COX-2 activity in cell culture. Notably, NTI-164 and CBD did not affect IL-5, IL-4 or IL-10 levels, which contrasts with previous studies demonstrating changes in the production of these cytokines induced by CBD under inflammatory conditions [93]. The variability in our findings and previous studies could be due to the differences in physiological responses to cannabis extracts, such as bell-shaped dose or dose-dependent responses [94], which needs to be explored in future studies. The neuroprotective and anti-inflammatory profiles of NTI-164 will be further characterised using *in vivo* models of neuroinflammation.

Although the underlying positive mechanisms of phytocannabinoids are challenging to characterise due to the complexity and diversity of the pharmacodynamic profile, a number of mechanisms for attenuation of inflammation have been proposed. A recent study showed that the anti-inflammatory effect of CBD involved inhibiting oxidative stress-activated NF-κB-dependent signalling pathway through regulation of NADPH production and glucose uptake [95]. In addition to the potential modulation of intracellular antioxidant pathways, other pharmacological anti-inflammatory mechanisms include modulating calcium signalling [96] and through increased peroxisome proliferator-activated receptor γ-dependent activation [97].

Apart from its anti-inflammatory effect, NTI-164 treatment significantly increased the proliferation of neural progenitors and promoted the survival of neurons under a cytotoxic state, whereas CBD had no effect. Congruent with our findings, CBD alone did not enhance neural progenitor cell proliferation [98], which could either indicate the observed effect was due to botanical synergy or due to variability of CBD dose-response. However, Luján and Valverde (2020) suggest that CBD exerts pro-neurogenic effects only after the generation of newborn neurons [99]. Nevertheless, the effect of phytocannabinoids on neural progenitor cell proliferation requires further investigation. Several biological mechanisms involved in CBD-mediated neuroplasticity protection and survival have been investigated as candidates [100], including extracellular-signal-regulated kinases [101], glycogen synthase kinase 3β signalling [102, 103], and mammalian target of rapamycin pathways [104]. Given the interaction between phytocannabinoids and the ECS, molecular mechanisms underlying the synergistic effect of cannabis constituents on neurogenesis in preclinical models need further investigation.

Together, these findings suggest that the anti-inflammatory effect of NTI-164 is likely due to the synergistic interaction of the cannabis extract derivatives rather than isolated CBD supporting the entourage effect of full-spectrum cannabis derivatives and suggesting a role for botanical synergy in the phenotypic transition of microglial cells. These results need to be confirmed and warrant further investigation of phytocannabinoid interaction in preclinical *in vivo* models to explore the potential for distinct effects of full-spectrum cannabis extracts compared to isolated compounds.

### Conclusions

Preclinical studies set the foundation for designing clinical trials using medicinal cannabis extract to treat various neuropathologies. In this study, the anti-inflammatory and neuroprotective effects of the chemically characterised full-spectrum cannabis extract NTI-164 were explored in immune-activated cells. NTI-164 showed higher efficacy than isolated CBD in attenuating inflammatory cytokines and promoting neuronal survival in the cytotoxic paradigm. Overall, these findings provide additional support for full-spectrum medicinal cannabis plant extract as a valid and safe therapeutic intervention for treating ASD-related and other neurological disorders. The growing evidence for the therapeutic benefit of whole-plant cannabis extracts in treating complex neurological disorders, which require a multi-mechanistic approach, is promising and supports further research. Future studies need to characterise the effects of NTI-164 on key anti-inflammatory and immunomodulatory mechanisms involved in neurological pathogenesis and progression *in vivo*. Also, the synergistic effect of phytocannabinoids in modulation of microglial cell functions warrants further investigation. Further examination of the optimal dose of NTI-164 and treatment duration may also enhance efficacy and expected outcomes.

## Acknowledgements

The supporting body played no role in the analysis or decision to publish this data.

